# Climate-linked biogeography of mycorrhizal fungal spore traits

**DOI:** 10.1101/2025.03.12.641256

**Authors:** Smriti Pehim Limbu, Sidney L. Stürmer, Geoffrey Zahn, Carlos A. Aguilar-Trigueros, Noah Rogers, V. Bala Chaudhary

**Affiliations:** Department of Environmental Studies, Dartmouth College, Hanover, NH 03755 USA; Universidade Regional de Blumenau, Departamento de Ciências Naturais, Blumenau, SC, 89030-903, Brazil; Biology Department, Utah Valley University, Orem, UT 84058 USA; Department of Biological and Environmental Science, University of Jyväskylä, Jyvaskyla, Finland

**Keywords:** spore traits, climate gradients, trait trade-offs, trait-environmental relationships, arbuscular mycorrhizal fungi, microbial traits

## Abstract

Climate-driven variation in microbial traits is crucial for predicting ecological responses to environmental change, yet global patterns remain understudied. Using global datasets of arbuscular mycorrhizal (AM) fungal observations linked to spore morphology, we show that climate gradients shape spore trait variation and functional diversity. Temperature and precipitation emerged as key drivers, influencing species range size and trait-environment relationships through trade-offs. Larger spore volumes were more prevalent in warm, wet, stable climates but were associated with smaller species range sizes, suggesting a trade-off between persistence and dispersal potential. Spores with ornamentation were also more prevalent in warm, wet climates and linked to restricted range sizes, possibly reflecting specialization to specific environmental conditions. Cell wall investment decreased in warmer, wetter climates, and was the strongest predictor of species range size, with intermediate investment associated with broader geographic distributions. Spore shape and color also exhibited climate-driven patterns, with spherical spores and greater pigmentation more common in warm, wet climates. Phylogenetic analyses revealed high trait conservatism for spore ornamentation, moderate for volume, low for color, and none for shape and cell wall investment. Additionally, functional diversity analyses revealed that warm, wet environments promote within-community trait richness but lower trait divergence, while broader climatic variability drives higher beta diversity. These findings highlight the role of climate in shaping microbial trait biogeography and suggest that evolutionary history constrains some traits while others are adaptable, suggesting that ongoing climate change may restructure AM fungal distributions, impacting plant-fungal interactions, nutrient cycling, and ecosystem stability.

**Significance statement:** A trait-based approach in microbial ecology helps explain how the environment shapes microbial traits, yet global patterns remain largely unknown. This study provides the first global assessment of climate influence on arbuscular mycorrhizal (AM) fungal spore traits. We identify key trade-offs between trait persistence and species range size, demonstrating that temperature and precipitation are primary drivers of spore volume, ornamentation, cell wall investment, shape, and color. These findings highlight the role of broad climatic patterns in shaping microbial communities and suggest that ongoing environmental change may alter AM fungal distributions, potentially disrupting plant-fungal interactions, soil health, and ecosystem stability.

## Introduction

As ecosystems undergo rapid environmental shifts, understanding species distributions and the ecological factors shaping them remains a critical challenge. Trait-based approaches, often hailed as the ‘Holy Grail of Ecology’ (1, 2), offer a unified framework for linking species traits to ecological functioning (3, 4) and community structure (5, 6), enabling predictions of species distributions and range shifts under climate change (7). Traits, defined as “measurable characteristics of an organism at the individual or other level of organization” (8), reveal ecological patterns by highlighting how specific traits confer advantages in different environments, leading to predictable trait-environment relationships (9). While much of this research has focused on plant traits (10–12), microbial traits are increasingly recognized for their role in predicting ecosystem functioning (13–15). However, major knowledge gaps remain regarding how microbial traits respond to climate drivers, limiting our ability to fully predict their ecological and evolutionary functions (16).

Microbes are fundamental to life and biogeochemical cycles (17), yet microbial traits remain understudied due to unique challenges. Unlike plants and animals, microbes exhibit vast genetic diversity, undergo horizontal gene transfer, and lack clearly defined individuals and species, complicating trait-based analyses (18). Paradoxically, these same challenges underscore the necessity of trait-based approaches, as taxonomy-based methods often fail to capture microbial ecological patterns and functional roles (18). Recent studies have highlighted how microbial trait variation is shaped by environmental factors (13, 16, 19). For instance, bacterial traits are correlated with soil pH, illustrating the role of environmental filtering in structuring microbial trait distributions (20). Given that microbial interactions span both aboveground and belowground systems, linking soil processes, plant health, and ecosystem stability (21), integrating trait-based approaches across these domains is essential for improving ecological modeling and conservation strategies.

Arbuscular mycorrhizal (AM) fungi provide an ideal model for studying microbial traits due to the widespread symbioses that they establish with approximately 72% of plant species (22) and their critical role in ecosystem functioning (23). These fungi enhance nutrient and water uptake, buffer plants against droughts and pathogens, and shape plant diversity and community structure (24–27). Additionally, AM fungi play a crucial role in global carbon, nitrogen, and phosphorus cycling (28). Their global distribution across diverse environments, from deserts to rainforests and boreal to tropical climates, underscores their adaptability. Although previous studies have improved our understanding of AM fungal distributions (29–31) and explored trait-environment relationships (32, 33), no global-scale trait-based analyses have been conducted for AM fungi or any other microbial groups to date. Given their obligate biotrophic lifestyle and high genetic diversity, trait-based methods are particularly valuable for understanding the ecological and functional roles of this group.

Variation in the traits of AM fungal spores – the reproductive and dispersal units of these fungi – reflects life history strategies and responses to climatic conditions across species (33–35). Key traits include size, shape, color, and ornamentation (projection or depression of the cell wall), which are used for morphological species identification. Several lines of evidence also highlight their role in dispersal (36, 37). AM fungal spore size ranges from 40 to 500 µm in diameter across species. This variation indicates distinct resource allocation patterns to reproduction across species: small-spored species produce more spores but invest less per spore, whereas large-spored taxa allocate more resources per spore, reflecting a trade-off between spore size and output (37). Although few studies have explicitly addressed the implications of size differences to dispersal, research on mushroom-forming fungi (the best studied fungal group in terms of traits) suggests that larger spores, endowed with greater nutrient reserves, tend to germinate under stressful conditions, whereas smaller spores, more likely to be wind-dispersed, remain dormant until favorable conditions arise (34, 38, 39). In these groups, smaller spores also travel greater geographical distances (40), potentially explaining differences in species range sizes (32). Spore wall thickness, which positively correlates with spore size (41), enhances resistance to chemical and microbial agents during passage through animal digestive tracts (42). Additionally, melanin – the pigment that gives dark colors to AM spores – has been found to increase with aridity (33), providing drought and fire resistance (35, 43). In spores of other fungal clades, melanin provides structural strength, UV protection, and resistance against harsh environmental conditions (39). Spore shape may influence surface area exposure to environmental conditions, with spherical shapes minimizing surface area relative to volume, thereby reducing exposure to desiccation and environmental stress (39). While the specific role of ornamentation in AM fungal spores remains unclear, it is speculated to aid in water repellence, dispersal, and substrate attachment, as observed in other fungal groups (39). Here, we analyzed multiple spore traits across diverse climates to determine how AM fungal species with distinct spore traits respond to global climate variation. Specifically, we examined five key traits – spore volume (a proxy for spore size), cell wall investment (spore wall thickness relative to spore volume), shape, color, and ornamentation – in relation to climate variables on a global scale.

Examining trait-environment relationships provides key insights into functional biogeography, defined by Violle et al. (44) as the patterns, causes, and consequences of trait diversity across geographic scales. This approach helps predict species responses to environmental change. However, analyzing traits individually may overlook trade-offs and interactions that shape community structure (45). A multi-trait approach captures these complexities by considering how traits respond to environmental gradients, interact with one another, and vary in importance under different conditions. This integrative perspective enhances predictions of microbial community dynamics and ecosystem functioning, overcoming the limitations of single-trait analyses.

We employed a data synthesis approach to investigate how microbial traits respond to environmental drivers, using AM fungal spore traits as a model system. We hypothesized that specific spore traits would correlate with temperature and precipitation patterns. Specifically, we expected (i) spore volume and cell wall investment to decrease in warmer, wetter conditions, as larger spores with thicker walls provide advantages in drier, cooler environments; (ii) ornamentation to be more prevalent in wetter regions, potentially aiding in water repellence; (iii) spores to become more spherical under low precipitation and temperature, minimizing surface area exposure; and (iv) melanin content of spore to increase with decreasing precipitation, enhancing drought resistance. We also hypothesized that trait alpha diversity would increase with temperature and precipitation, while beta diversity would rise with greater environmental heterogeneity due to distinct selection pressures. Therefore, we predicted that (v) AM fungal spore traits influence species’ geographic distribution range by mediating tolerance to climate and biome conditions, and (vi) trait-environment relationships would be phylogenetically structured, with closely related species exhibiting similar adaptations due to shared evolutionary histories, indicating evolutionary constraints in shaping AM fungal responses to climate gradients. By integrating climate-trait relationships on a global scale, our study provides a comprehensive framework for understanding AM fungal biogeography and offers a scalable model for predicting how these communities will respond to environmental change across diverse ecosystems.

## Results

### Climate gradients shape community-weighted AM fungal spore traits

The first four principal components (PCs) from the Principal Component Analysis (PCA) of climate variables explained approximately 89% of the total environmental variation (Fig. S1). PC1, here referred to as the “Temperature and Moisture Gradient,” captured the combined variation in mean annual temperature and precipitation. PC2, here referred to as the “Precipitation Variability Gradient,” represented the variation in annual precipitation seasonality. Similarly, PC3 captured precipitation extremes, and PC4 captured the variation in soil pH and annual wind gradient. The “Temperature and Moisture Gradient” emerged as a significant predictor of all five traits (Table 1; Fig. 2). Additionally, biome, target gene, and spatial coordinates (latitude and longitude) also consistently explained substantial variation across all traits (*P < 0.001*), emphasizing the importance of ecological context, and geographic variation (Table 1). The spatial effect demonstrated non-uniform trait distributions across regions, with certain geographic areas exhibiting stronger environmental filtering effects than others (Fig. S5). Furthermore, the year of sampling was also a significant random effect for spore volume, investment, shape and color (*P < 0.001*), indicating that both spatial and temporal variations influence trait patterns.

**Table 1.**
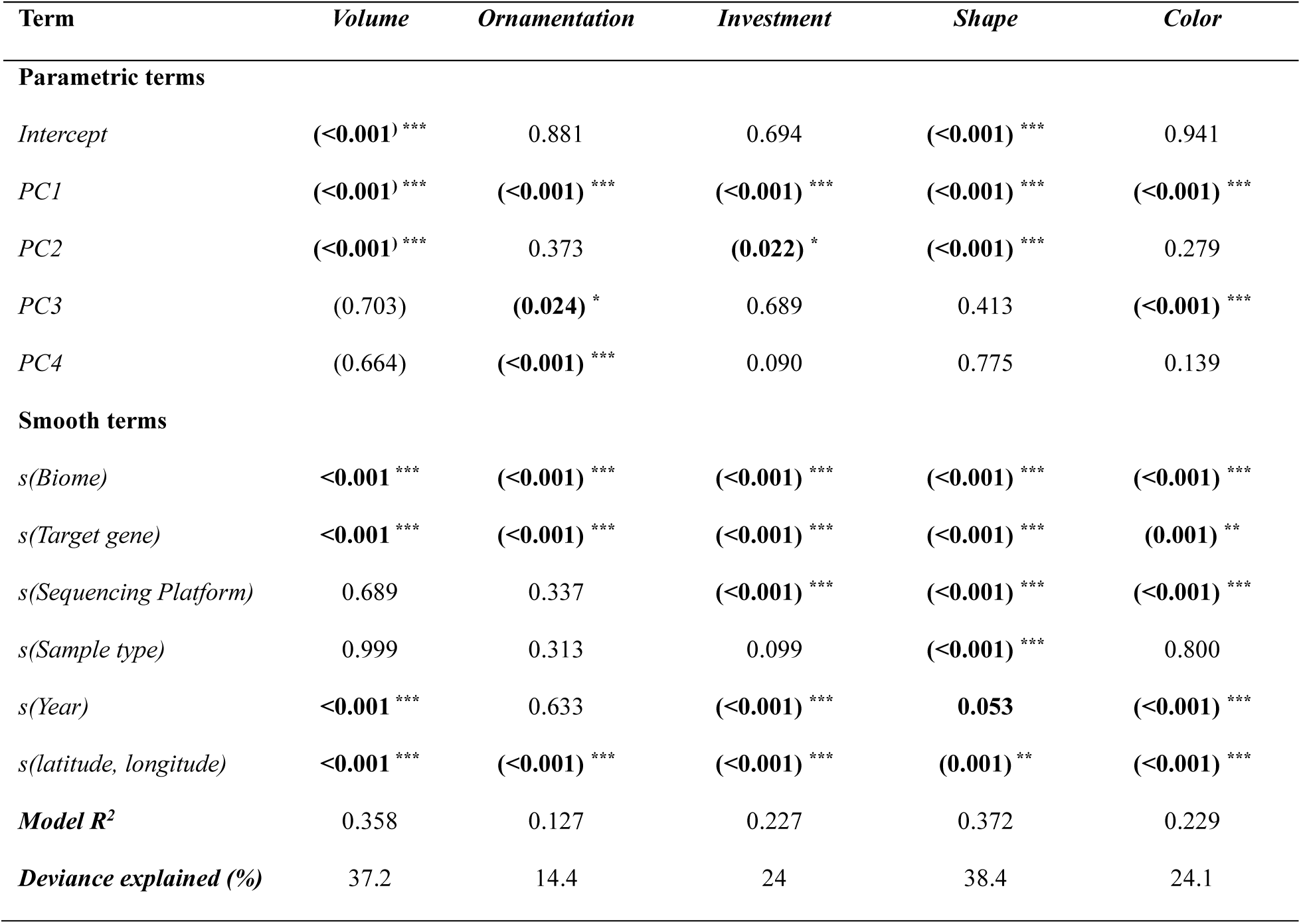
Relationship between the community-weighted AM fungal spore traits and environmental variables. The table presents estimates P-values for parametric and smooth terms, including the first four principal components (PC1, PC2, PC3, and PC4), which capture gradients in temperature, precipitation, soil pH, and wind patterns across sampling sites. The final rows include model R² and the percentage of deviance explained, indicating model fit for each trait. Significance levels are indicated as follows: * for *P* < 0.05, ** for *P* < 0.01, and *** for *P* < 0.001.

| Term | Volume | Ornamentation | Investment | Shape | Color |
| --- | --- | --- | --- | --- | --- |
| **Parametric terms** |  |  |  |  |  |
| *Intercept* | (<0.001) *** | 0.881 | 0.694 | (<0.001) *** | 0.941 |
| *PC1* | (<0.001) *** | (<0.001) *** | (<0.001) *** | (<0.001) *** | (<0.001) *** |
| *PC2* | (<0.001) *** | 0.373 | (0.022) * | (<0.001) *** | 0.279 |
| *PC3* | (0.703) | (0.024) * | 0.689 | 0.413 | (<0.001) *** |
| *PC4* | (0.664) | (<0.001) *** | 0.090 | 0.775 | 0.139 |
| **Smooth terms** |  |  |  |  |  |
| *s(Biome)* | <0.001 *** | (<0.001) *** | (<0.001) *** | (<0.001) *** | (<0.001) *** |
| *s(Target gene)* | <0.001 *** | (<0.001) *** | (<0.001) *** | (<0.001) *** | (0.001) ** |
| *s(Sequencing Platform)* | 0.689 | 0.337 | (<0.001) *** | (<0.001) *** | (<0.001) *** |
| *s(Sample type)* | 0.999 | 0.313 | 0.099 | (<0.001) *** | 0.800 |
| *s(Year)* | <0.001 *** | 0.633 | (<0.001) *** | 0.053 | (<0.001) *** |
| *s(latitude, longitude)* | <0.001 *** | (<0.001) *** | (<0.001) *** | (0.001) ** | (<0.001) *** |
| **Model $R^2$** | 0.358 | 0.127 | 0.227 | 0.372 | 0.229 |
| **Deviance explained (%)** | 37.2 | 14.4 | 24 | 38.4 | 24.1 |

**Figure 1.**
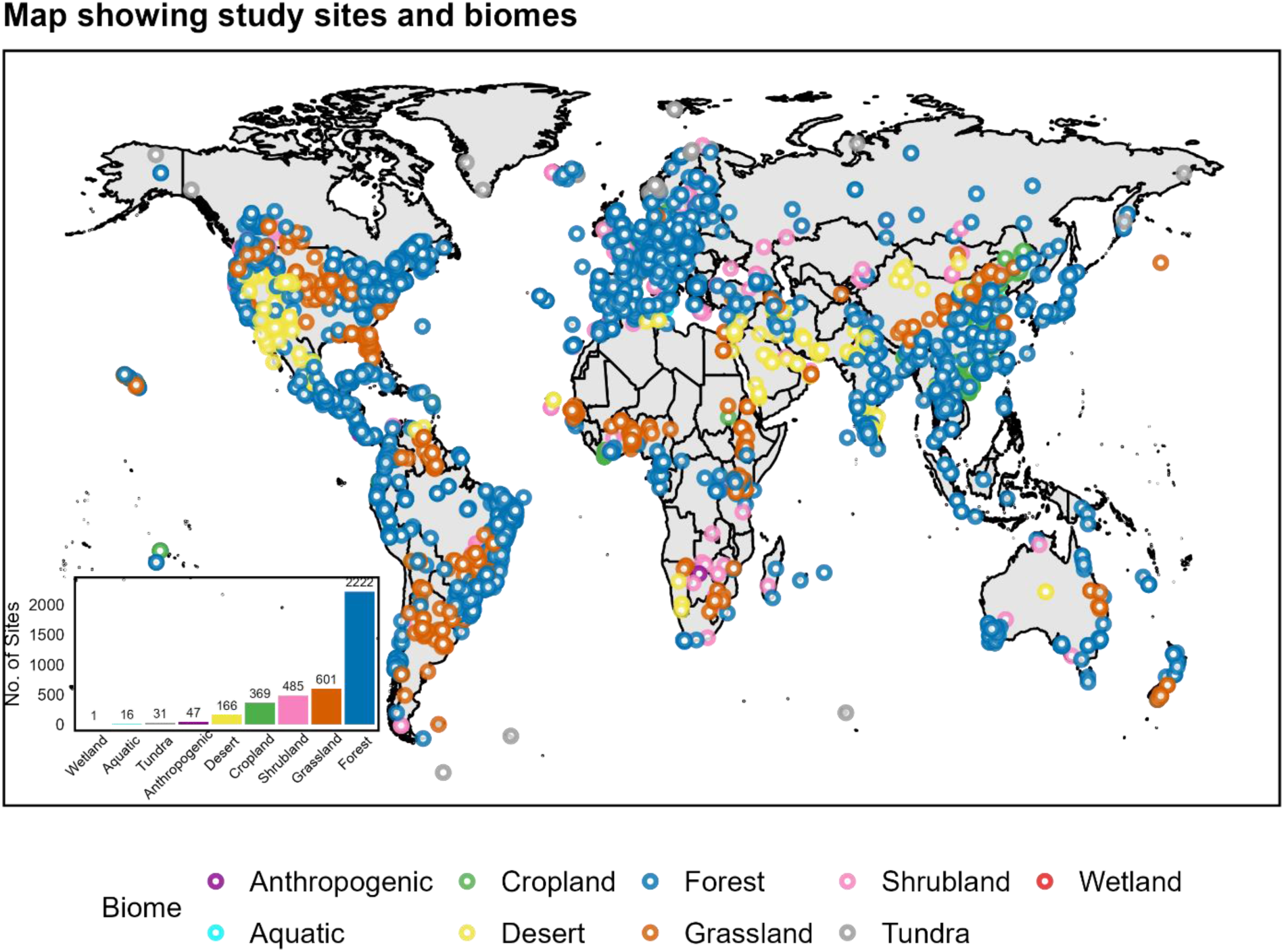
Global distribution of study sites used in the analysis of arbuscular mycorrhizal (AM) fungal spore traits across diverse biomes. The inset bar plot shows the number of study sites per biome, with ‘Forest’ being the most represented (N = 2,222 sites) and ‘Wetland’ the least (N = 1 site). Each biome is color-coded as indicated in the legend, corresponding to its respective sites on the map.

**Figure 2.**
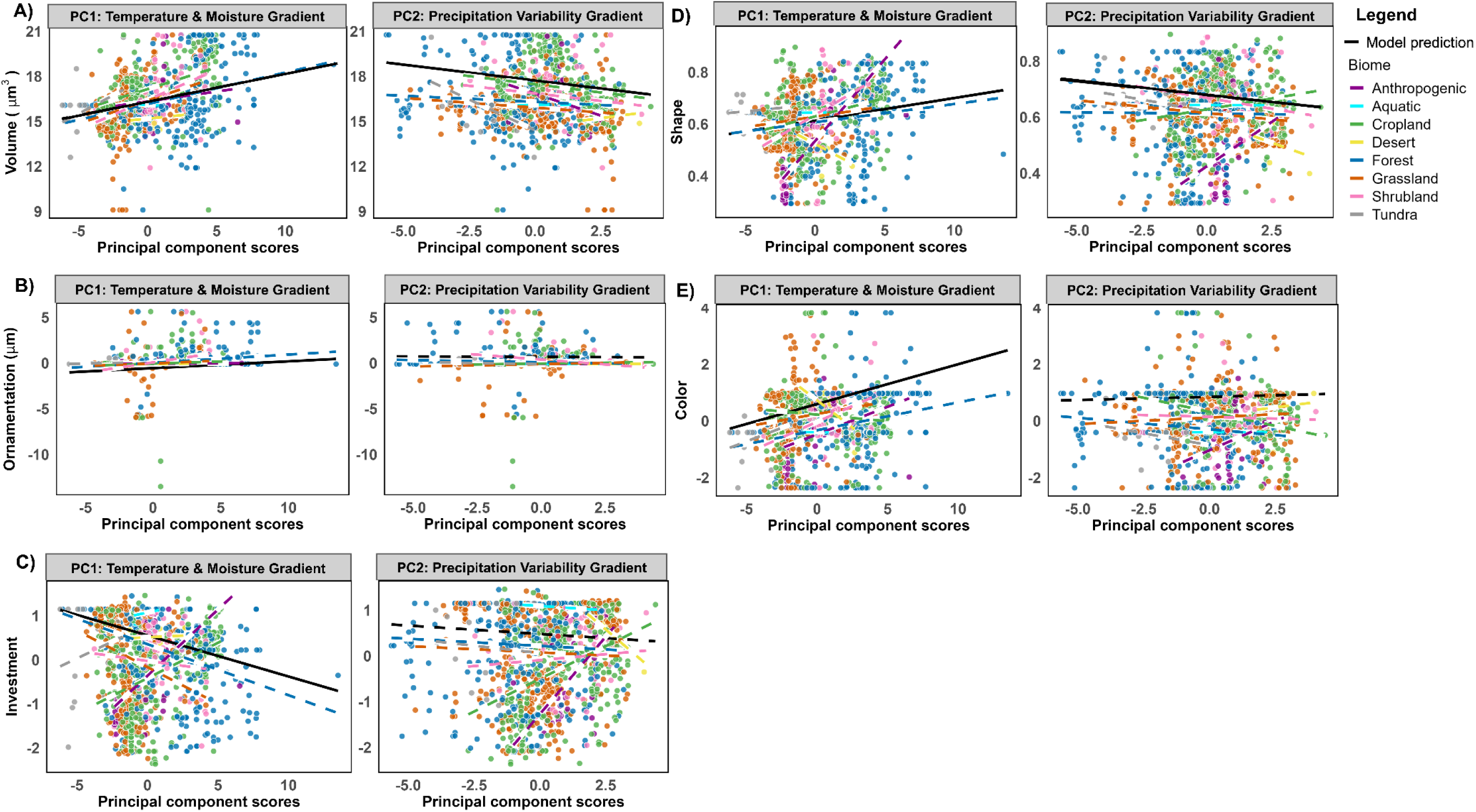
Relationship between environmental gradients and arbuscular mycorrhizal (AM) fungal spore traits across biomes. Panels (A–E) depict the effects of PC1 (Temperature & Moisture Gradient; left column) and PC2 (Precipitation Variability Gradient; right column) on spore traits: (A) Volume, (B) Ornamentation, (C) Cell wall investment, (D) Shape, and (E) Color. Traits were transformed to meet the normality assumptions of the model: Volume and Ornamentation were Box-Cox transformed, while Investment, Shape, and Color were Yeo-Johnson-transformed. Data points represent observed trait values, color-coded by biome type. Solid black lines indicate significant model (Generalized Additive Model) predictions (P < 0.05), while dashed black lines indicate non-significant predictions. Colored dashed lines represent biome-specific linear trends.

Spore volume (Fig.2A), ornamentation (Fig. 2B), shape (Fig. 2D) and color (Fig.2E) showed significant positive associations with the “Temperature and Moisture Gradient,” suggesting that higher temperatures and moisture favor communities with larger spore volume, greater ornamentation, more spherical shapes, and darker pigmentation. In contrast, spore cell wall investment (Fig. 2C) was negatively associated with the “Temperature and Moisture Gradient,” indicating that warmer and wetter conditions favor communities with reduced cell wall investment.

PC2, the “Precipitation Variability Gradient,” had weaker effects on most traits compared to PC1. Spore volume and shape showed significant negative associations with PC2 (*P < 0.001*), while ornamentation, investment, and color displayed no significant patterns (Table 1). Spore ornamentation was negatively associated with PC3 (precipitation extremes) (*P = 0.024*), whereas spore color exhibited a significant positive association with this gradient (*P < 0.001*). Additionally, spore ornamentation demonstrated a significant positive relationship with PC4 (soil pH and annual wind gradient) (*P < 0.001*).

### Environmental variables predict functional alpha diversity

The first four PCs (see above) from the PCA of climate variables explained approximately 90% of the total environmental variation in functional diversity metrics (Fig. S3). Functional richness exhibited a significant positive association with the “Temperature and Moisture Gradient” (*P < 0.001*), indicating that sites with higher temperatures and moisture levels tend to support greater functional richness (Table 2; Fig. 3A). Additionally, functional richness displayed a weaker positive relationship with the “Precipitation Variability Gradient” (*P = 0.02*). In contrast, functional evenness showed no significant trends with either PC1 or PC2, suggesting that the uniformity of trait distribution within sites is not influenced by temperature, moisture, or precipitation variability gradients (Fig. 3B). Functional divergence, however, exhibited a significant negative association with the “Temperature and Moisture Gradient” (*P = 0.006*), implying that higher temperature and moisture levels reduce trait divergence of AM fungal communities within sites (Fig. 3C). No significant patterns were observed between functional divergence and PC2.

**Table 2.** Relationship between the functional diversity of AM fungal spore traits and environmental variables. The table presents estimates P-values for parametric and smooth terms, including the first four principal components (PC1, PC2, PC3, and PC4), which capture gradients in temperature, precipitation, soil pH, and wind patterns across sampling sites. The final rows include model R² and the percentage of deviance explained, indicating model fit for each trait. Significance levels are indicated as follows: * for *P* < 0.05, ** for *P* < 0.01, and *** for *P* < 0.001.

| Term | Functional richness | Functional evenness | Functional divergence |
| --- | --- | --- | --- |
| **Parametric terms** |  |  |  |
| *Intercept* | 0.066 | (**<0.001**) *** | (**<0.001**) *** |
| *PC1* | (**0.001**) ** | 0.619 | (**0.006**) ** |
| *PC2* | **0.023** * | 0.981 | 0.667 |
| *PC3* | 0.168 | 0.127 | 0.368 |
| *PC4* | 0.178 | 0.913 | 0.385 |
| **Smooth terms** |  |  |  |
| *s(Biome)* | 0.422 | (**<0.001**) *** | **0.031** * |
| *s(Target gene)* | (**<0.001**) *** | (**0.005**) ** | (**<0.001**) *** |
| *s(Sequencing Platform)* | (**<0.001**) *** | 0.399 | (**0.040**) * |
| *s(Sample type)* | (**<0.001**) *** | **0.013** * | (**0.001**) ** |
| *s(Year)* | (**<0.001**) *** | 0.378 | 0.391 |
| *s(latitude, longitude)* | (**<0.001**) *** | **0.036** * | (**<0.001**) *** |
| **Model $R^2$** | 0.448 | 0.463 | 0.561 |
| **Deviance explained (%)** | 48.3 | 52 | 61 |

**Figure 3.**
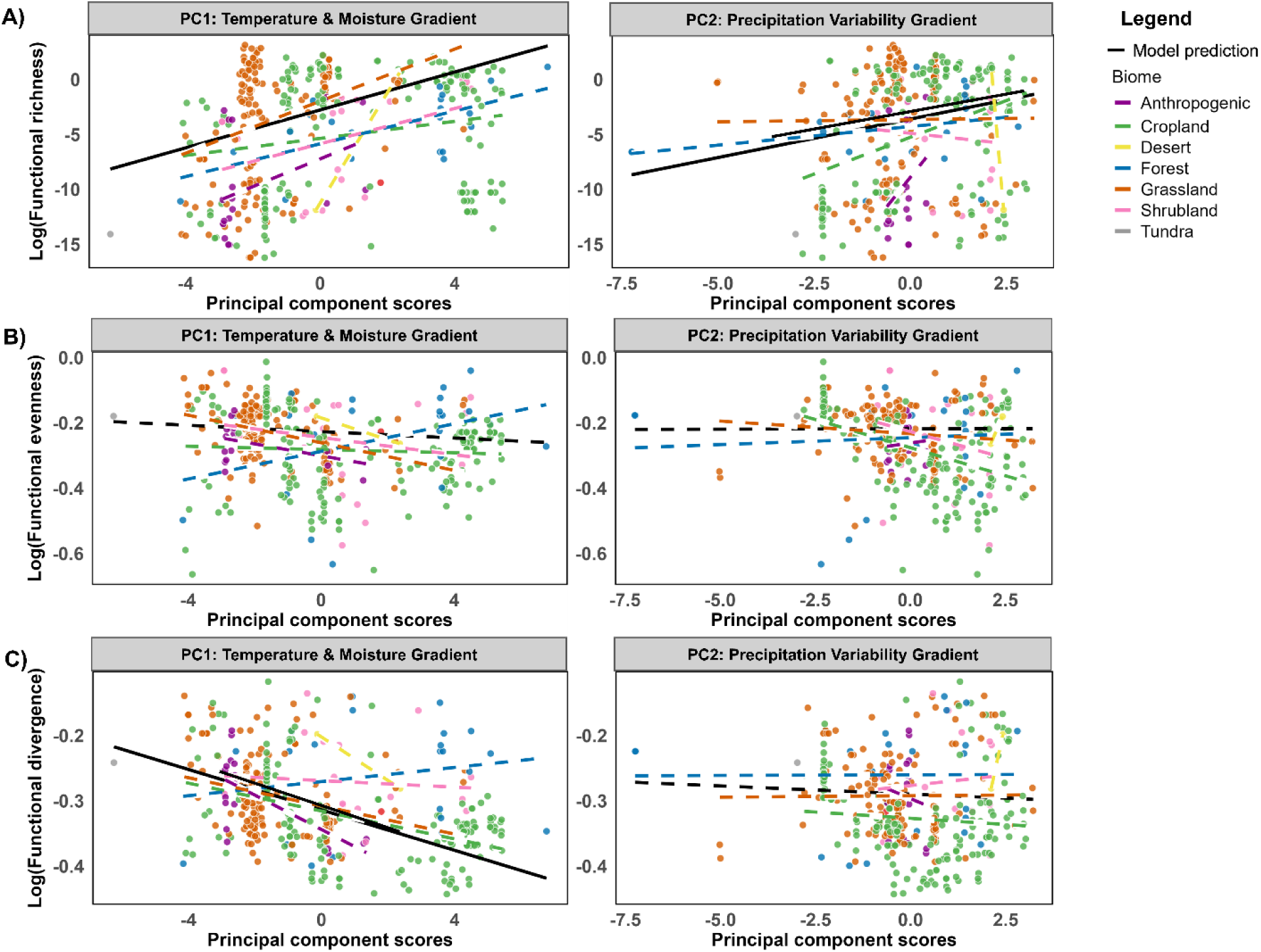
Relationship between functional alpha diversity indices. A: Functional Richness, B: Functional Evenness, C: Functional Divergence) and the first two Principal Components (PCs) derived from environmental variables across different biomes (N = 1,248). The left column represents associations with PC1 (Temperature & Moisture Gradient), while the right column represents associations with PC2 (Precipitation Variability Gradient). Each point represents a site, color-coded by biome type as indicated in the legend. Smooth dashed lines depict fitted relationships for each biome, highlighting biome-specific responses to environmental gradients. The black line represents the overall trend across biomes predicted by generalized additive models, with solid lines indicating significant relationships (P < 0.05) and dashed lines representing non-significant relationships. Functional diversity indices were log-transformed to meet model assumptions.

### Environmental variables drive trait beta-diversity

Generalized Dissimilarity Modeling (GDM) revealed that environmental variables strongly influence the beta diversity of AM fungal spore traits. Predicted trait dissimilarity, calculated as the Euclidean distance among traits, showed a clear positive association with observed environmental dissimilarity, indicating that sites with greater environmental differences exhibited higher divergence in AM fungal spore traits (Fig. 4A). The strong relationship between predicted and observed dissimilarity demonstrated a good model fit, effectively capturing trait-environment relationships (Figure 4B). Mean annual precipitation showed a linear increase in trait dissimilarity with increasing precipitation, while soil pH exhibited a steep initial rise in trait dissimilarity up to soil pH 6, stabilizing at higher pH levels (Fig. 4C-D). Mean annual temperature displayed a gradual, nonlinear increase in dissimilarity across the temperature gradient, indicating a progressive trait response to thermal variation (Fig. 4E). Precipitation seasonality revealed a sharp initial increase in trait dissimilarity that plateaued at higher seasonality values, suggesting a threshold effect of precipitation variability on trait dissimilarity (Fig. 4F).

**Figure 4.**
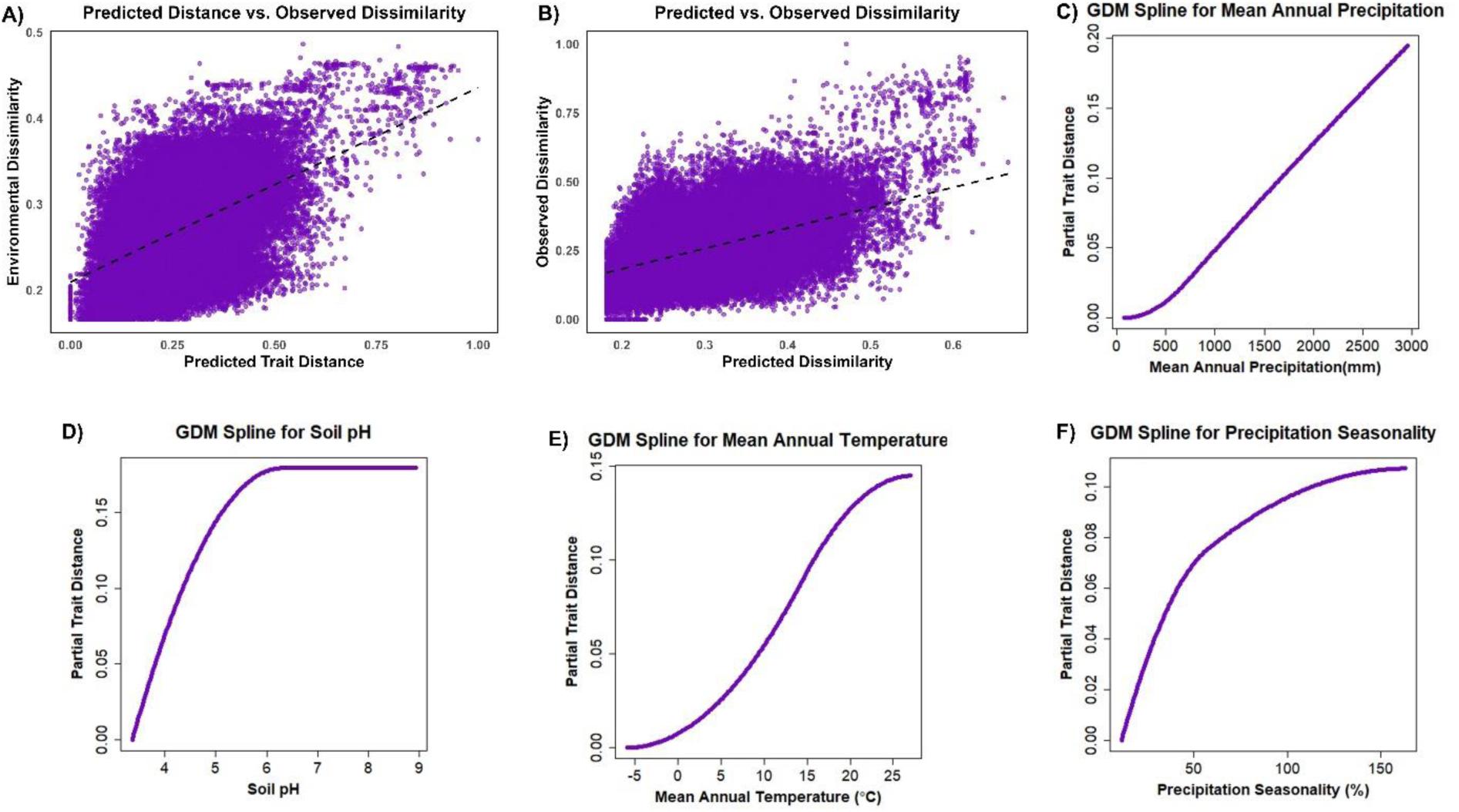
Generalized Dissimilarity Modeling (GDM) results showing functional beta diversity of AM fungal spore traits (volume, ornamentation, cell wall investment, shape, and color. (A) Positive relationship between predicted trait distance (Euclidean distance of traits across unique sites) and observed environmental dissimilarity. (B) Model fit shown by the relationship between predicted dissimilarity and observed dissimilarity, with a linear trend indicated by the dashed line. (C-F) GDM splines illustrate the contributions of key environmental predictors to trait dissimilarity: (C) *Mean Annual Precipitation (mm)*: Partial trait distance increases linearly with precipitation. (D) *Soil pH*: Partial trait distance rises steeply at lower pH and plateaus at higher pH. (E) *Mean Annual Temperature (°C)*: Nonlinear increase in partial trait distance along the temperature gradient. (F) *Precipitation Seasonality (%)*: Sharp initial rise in partial trait distance followed by stabilization. Spline plots (C-F) show the functional relationships between environmental predictors and trait dissimilarity, measured as partial trait distance.

### Phylogenetic history conserves some spore traits while others vary

Arbuscular Mycorrhizal fungal spore traits exhibited varying degrees of phylogenetic signal. Pagel’s λ values for spore traits such as ornamentation (λ = 0.78), volume (λ = 0.41), color (λ = 0.19) indicate strong, moderate, and low phylogenetic signals, respectively (Table 3), suggesting that these traits are at least partially conserved across evolutionary lineages. In contrast, spore shape and cell wall investment exhibited negligible phylogenetic signal (λ ≈ 0; Table 3), reflecting substantial variability largely independent of phylogenetic constraints. Certain phylogenetic groups demonstrated conservation of traits, while others displayed trait divergence (Fig. 5).

**Table 3.** Phylogenetic signal of AM fungal spore traits estimated using Pagel’s Lambda (λ). Higher λ values indicate stronger phylogenetic conservation, suggesting that closely related species share similar trait values. Conversely, λ values closer to zero indicate minimal phylogenetic signal, reflecting substantial trait variability independent of phylogenetic constraints.

| Trait | Estimated Pagel's Lambda ($\lambda$) | Log-Likelihood |
| --- | --- | --- |
| Volume | 0.412561 | -3235.82 |
| Ornamentation | 0.781278 | -365.974 |
| Cell wall investment | 7.33137e-05 | 71.0581 |
| Shape | 7.33137e-05 | 146.638 |
| Color | 0.192061 | -368.557 |

**Figure 5:**
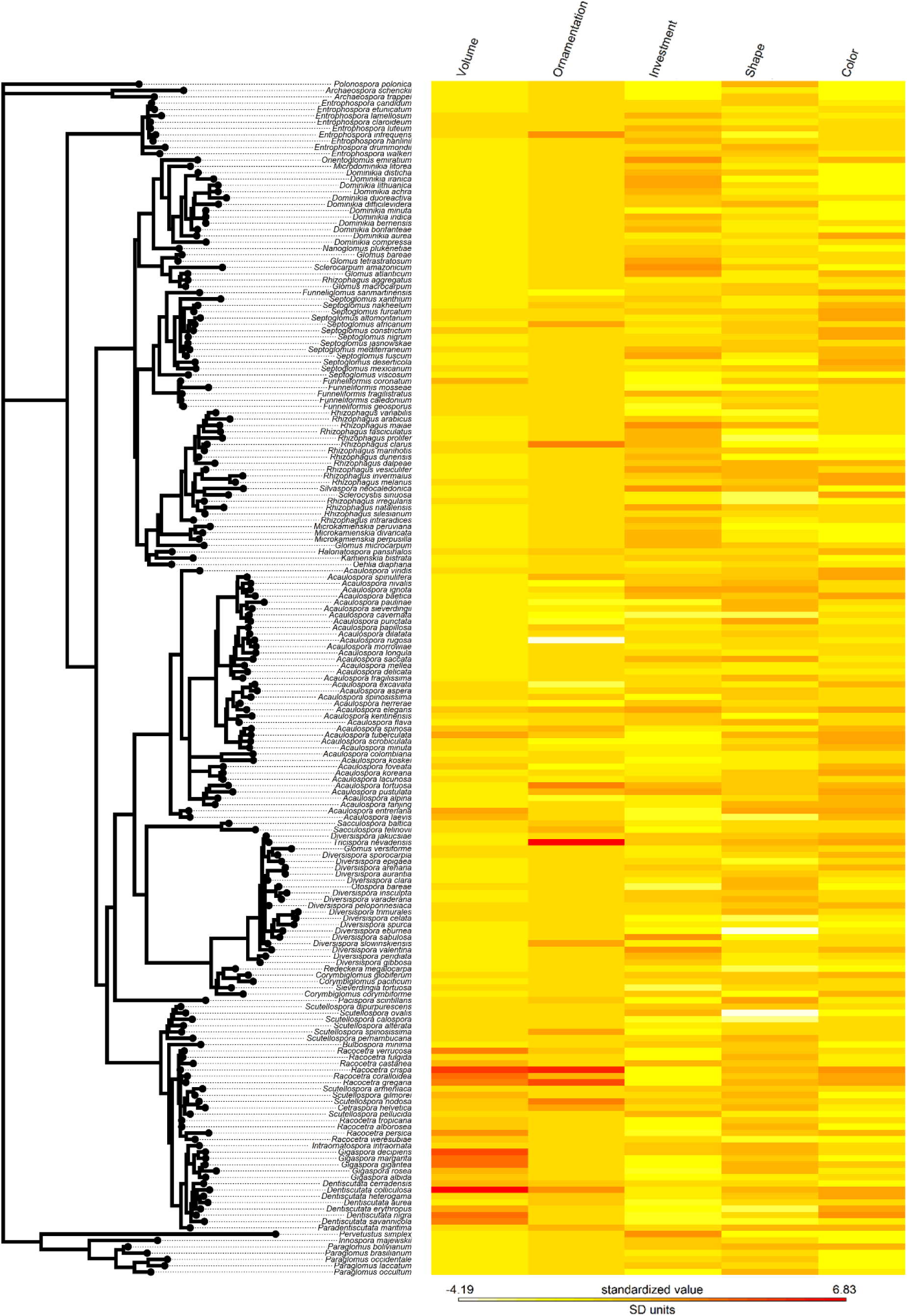
Phylogenetic heatmap of AM fungal spore traits. The phylogenetic tree (left) represents the evolutionary relationships among arbuscular mycorrhizal (AM) fungal species. The heatmap (right) displays standardized values of five key spore traits: volume, ornamentation, cell wall investment, shape, and color across species. Warmer colors (e.g., red) indicate higher trait values, while cooler colors (e.g., yellow) represent lower trait values. The combination of phylogenetic relationships and trait variation highlights the degree of phylogenetic conservatism and divergence in AM fungal spore traits, showcasing the interplay between evolutionary history and trait adaptations.

### Species range size is affected by traits

The random forest model identified spore traits as the primary predictors of AM fungal species range size. Among all the predictors, spore cell wall investment emerged as the most important predictor, followed by color, shape, and volume (Fig. 6A). Environmental variables such as mean annual temperature and mean annual precipitation also contributed to the model but were less influential compared to the spore traits (Fig. 6A). Variables like soil pH, mean annual wind and precipitation seasonality exhibited minimal contributions. These results highlight the dominant role of spore traits over the measured environmental variables in shaping species’ range sizes.

**Figure 6:**
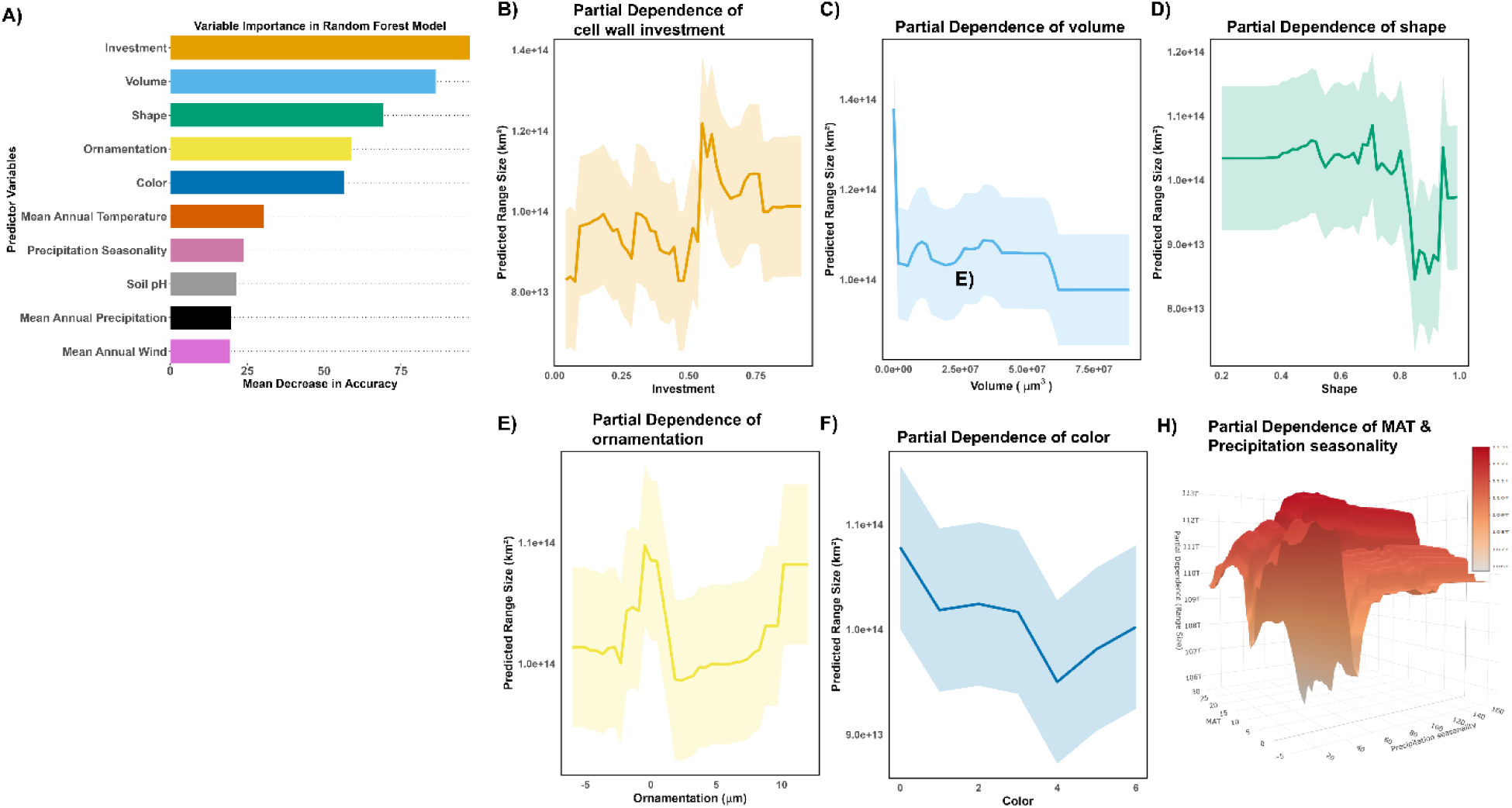
Random Forest model results illustrating the influence of AM fungal spore traits and environmental variables on species range size. (A) Variable importance plot showing the relative contribution of predictor variables to species range size, measured as Mean Decrease in Accuracy in the Random Forest model. Spore traits (cell wall investment, volume, shape, ornamentation, and color) had the highest predictive importance, while environmental variables (mean annual temperature, precipitation seasonality, soil pH, and wind) had comparatively lower contributions. (B–F) Partial dependence plots illustrating the relationship between individual spore traits and predicted species range size (km²) (G) predicted species range size based on interactive effect of mean annual temperature (MAT) and precipitation seasonality, with elevation and color scale representing species range size across different trait values.

## Discussion

Leveraging a harmonized global dataset of AM fungal spore traits, our study underscores the pivotal role of climate in shaping microbial trait distributions and functional diversity. Temperature has long been recognized as a key climatic factor influencing fungal species distribution (46), and as the dominant abiotic driver of AM fungal community structure at regional scales (47). In this study, temperature and moisture gradient and precipitation variability emerged as primary environmental drivers of trait variation, with all five AM fungal spore traits showing significant correlations with these gradients. These findings reinforce the role of environmental filtering in structuring AM fungal communities and support our initial hypothesis. However, climate-driven trade-offs were evident, where traits ecologically advantageous under specific climate conditions, simultaneously limited species range sizes. Spore cell wall investment and volume were strong predictors of species range sizes, emphasizing their significance in determining geographic distributions. Additionally, phylogenetic analyses revealed strong conservatism in spore ornamentation, moderate conservatism in spore volume, and low conservation in color, whereas shape, and cell wall investment showed no conservatism, indicating stronger influence of local environmental conditions. The interplay between environmental filtering, evolutionary history, and trait variation shapes AM fungal community structure, ultimately influencing species range sizes and geographic distributions. As climate change continues to alter temperature and precipitation patterns, AM fungal distributions are likely to shift, potentially impacting plant-AM fungal symbioses, nutrient cycling, and ecosystem stability.

### Trade-offs and climate-driven patterns in AM fungal spore traits and species range sizes

Contrary to expectations, higher temperatures and precipitation promoted communities with larger spore volume, while greater precipitation seasonality favored communities with smaller spore volume. This suggests that larger spores provide adaptive advantages in warmer, wetter climates with lower precipitation variability, as their greater nutrient reserves enhance spore germination and establishment (39), particularly during the asymbiotic phase where AM external hypha rely on spore reserves to grow and contact host roots (48). Consistent resource availability in warm and wet climates may allow species with larger spores to optimize resource use and outcompete smaller-spored species. In contrast, high precipitation variability may favor species producing smaller spores due to lower resource investment and greater dispersal potential, facilitating rapid colonization in unpredictable environments (49, 50). A previous study has showed that some species in the Gigasporaceae family exhibited variable spore volumes along temperature gradients in sand dunes, but these correlations were not significant (47).

As also noted by Chaudhary et al. (34), larger spores were associated with reduced species range sizes, suggesting a trade-off between spore size and dispersal potential: while increased spore volume enhances survival success in stable, resource-rich conditions, it likely limits dispersal, thereby restricting geographic range. Similarly, in North American ecoregions, wind-dispersed were exclusively small-spored *Glomus* (51), suggesting that spore dispersion is influenced by interaction between spore traits and environmental conditions (52). In contrast, Kivlin (53) found no correlation between AM fungal spore size and range size after accounting for phylogenetic relationships, suggesting that shared evolutionary history may obscure ecological patterns. In our study, we did not apply phylogenetic corrections for two reasons: first, our objective was to investigate the ecological relationships between spore traits and environmental gradients without partitioning out the phylogenetic signal, as this could mask adaptive responses to current environmental conditions (54); second, our phylogenetic tree contained fewer species, which would reduce statistical power and potentially bias phylogenetic comparative analyses. Despite this approach, moderate phylogenetic conservatism observed for spore volume suggests that evolutionary history plays a key role in shaping this trait (55). The evolutionary persistence of larger spore sizes may be linked to higher germination success and stress tolerance (37). However, these evolutionary constraints may limit the adaptability of AM fungal spore traits to changing climate.

Presence of ornamentation in AM fungal spores increased with higher temperatures, moisture levels, and wind gradients, supporting its potential roles in water repellence and dispersal (39, 56). This finding aligns with our prediction that ornamented spores would be more prevalent in wetter environments, where different types of ornamentation, such as surface projections (warts, papillae) or depressions (pits), may improve spore stability by repelling water. Additionally these structures could enhance dispersal potential by reducing clumping in humid conditions, facilitating efficient transport through water or wind (57).

Our results revealed a strong phylogenetic signal for ornamentation, indicating that this trait is highly conserved within evolutionary lineages. This pattern suggests that ornamentation is not solely shaped by environmental filtering but is also constrained by evolutionary history. One possible explanation of phylogenetic conservatism could be that ornamentation provides consistent adaptive advantages, such as enhanced protection against UV radiation, desiccation, and microbial attack, leading to its retention within lineages (56) However, despite this evolutionary conservatism, spores lacking ornamentation were associated with larger range sizes. This pattern suggests a trade-off between localized dispersal potential and broader ecological flexibility. Specifically, while ornamentation enhances localized dispersal by stabilizing spores in high-wind conditions, it may also increase habitat specialization, restricting species to narrower ecological niches. In contrast, smoother spores, which lack specialized surface structures, may experience less drag during dispersal and exhibit greater ecological adaptability, enabling colonization across diverse habitats (58).

Like spore volume, spores will lower cell wall investment were more prevalent with increasing temperature and moisture gradient. While fungal cell walls primarily provide structural reinforcement and protection, they also regulate spore transitions between developmental stages, such as germination and dormancy (59). Lower investment in cell wall thickness in warmer, wetter conditions may facilitate germination by reducing energy costs for structural maintenance (60). Additionally, cell wall investment was the strongest predictor of species range size, with intermediate investment correlating with the broadest geographic distributions. This pattern suggests a trade-off between structural protection and dispersal, where excessive investment may limit dispersal, while minimal investment could compromise spore survival in variable environments.

Contrary to our prediction, more spherical spores were more common than less spherical spores with increasing temperature and moisture, while less spherical spores were more common with greater precipitation variability. In stable, warm, and wet environments, consistent moisture likely reduces selective pressure for elongated or irregular spore shapes, which aid in attachment and retention in fluctuating climatic conditions (61). Conversely, in regions with high precipitation variability, deviations from sphericity may enhance spore anchoring to substrates (39). Additionally, both elongated and perfectly spherical spores were associated with broader geographic ranges than nearly spherical spores, suggesting a nonlinear relationship between shape and dispersal potential. Perfect spheres may facilitate passive transport via wind or water, while intermediate shapes may improve substrate attachment and localized establishment. In contrast, nearly spherical spores may face dispersal inefficiencies, limiting their geographic range.

Under warmer and wetter conditions, AM fungal communities were more likely to be composed of darker spored taxa, potentially reflecting a protective role of pigmentation against UV radiation and desiccation (33). Pigmentation can also provide resistance to heat and other stresses associated with fire (62). However, species range size increased with lower pigmentation, suggesting that lighter-colored spores may be more adaptable across diverse environments. The low phylogenetic conservatism observed for spore color suggests weak evolutionary constraints on this trait, with pigmentation variability conserved within certain lineages. For instance, species of *Septoglomus* predominantly produce dark-colored spores while those of *Dominikia* typically have light-colored (hyaline to creamy) spores. Spore color has only recently been recognized as a trait that responds to environmental conditions. Hopkins et al. (62) demonstrated that spore color luminance decreased after a fire event for *Gigaspora spp.*, *Cetraspora pellucida* and *Paraglomus spp.* in a tall grass prairie restoration area, while spore color saturation was highly variable among AM fungal species in response to fire. Additionally, AM fungal spore luminance was not significantly different along an aridity gradient but melanin tended to increase with aridity (33). Together, these findings reveal intricate relationships between AM fungal spore traits, environmental gradients, and evolutionary history. The trade-offs observed in spore traits highlight the balance between survival strategies, dispersal efficiency, and environmental filtering across biomes.

### Functional diversity: Alpha and beta diversity patterns

By analyzing multiple traits simultaneously, our study provides new insights into how climate shapes the functional diversity of AM fungal spores. Functional richness increased with higher temperatures and moisture levels, suggesting that warm, wet environments enhance resource availability and promote the coexistence of species with diverse traits (63). In contrast, functional evenness showed no significant relationship with climate, suggesting that biotic interactions, rather than climate, may regulate trait distribution within communities, as observed in plant communities (64). However, functional divergence declined with temperature and moisture gradient, suggesting that warm, wet conditions favor a narrower range of traits due to competitive filtering, which selects for species with similar ecological strategies (65).

At the regional scale, beta diversity increased with environmental heterogeneity, highlighting the role of climatic variability in shaping AM fungal spore trait differentiation across sites. Temperature, precipitation, and soil pH emerged as key drivers of beta diversity, likely shaping distinct environmental regimes that select for specific trait adaptations. For example, AM fungal communities in arid environments may favor drought-tolerant traits, whereas those in wetter, stable climates may prioritize traits linked to continuous resource acquisition (66, 67). This pattern illustrates how spatial environmental variability fosters distinct trait assemblages; even as environmental filtering reduces local trait divergence within communities.

Our study demonstrates how broad-scale climate gradients shape AM fungal functional diversity across local and regional scales. At the local scale, environmental filtering may favor specific traits, leading to reduced trait variability within communities. For instance, abiotic factors such as soil pH and nutrient availability act as key filters, selecting for AM fungal taxa adapted to these conditions (68). At the regional scale, climatic variability differentiates trait composition among communities, contributing to higher beta diversity in regions with greater environmental heterogeneity. These results support the climate-filtering hypothesis (69), where environmental conditions shape AM fungal trait distributions through selective pressures operating at multiple spatial scales.

### Limitations

While our study provides valuable insights into how climate gradients shape AM fungal spore traits, several limitations must be considered. First, our data were synthesized from multiple sources, and despite efforts to standardize methodologies, potential biases may arise due to inconsistencies in sampling techniques, sequencing platforms, and geographic coverage, particularly in underrepresented biomes. Second, we did not account for intraspecific trait variation, which could provide deeper insights into trait-environment relationships by capturing within-species plasticity. Third, our study focused on a subset of AM fungal spore traits, potentially overlooking other relevant traits that influence ecological interactions and adaptation.

Additionally, our study primarily examined acaulosporoid spores, potentially weakening environmental signals if dimorphic taxa exhibited a dispersal vs. persistence trade-off by changing spore types in response to environmental conditions. This is particularly relevant because dimorphic spores may shift between dispersal-optimized and persistence-optimized forms, masking environmental influences on spore traits. However, this limitation also strengthens our findings; the clear trait-climate patterns observed suggest that environmental filtering is robust even without considering glomoid spore forms. If dimorphic taxa indeed alter spore types due to environmental trade-offs, the observed patterns would be even stronger.

Lastly, while we identified significant associations between traits and climate gradients, the causal mechanisms remain speculative and require validation through controlled experiments. Future research integrating manipulative experiments, high-resolution trait data, and expanded trait assessments will be essential for refining our understanding of AM fungal functional biogeography.

### Implications and future directions

Despite limitations, our findings have important ecological and practical implications. The global scope of this study underscores the pervasive influence of climate gradients on microbial functional diversity, emphasizing the vulnerability of AM fungal communities to climate change. Understanding these trait-environment relationships is crucial for predicting shifts in AM fungal distributions and their cascading effects on plant communities, nutrient cycling, and ecosystem stability. Furthermore, our study reinforces the value of trait-based approaches in microbial ecology, providing a framework for guiding conservation strategies and ecosystem management in a rapidly changing world. By linking microbial traits to environmental drivers, this approach enhances our ability to forecast community responses to climate change. Future research should integrate experimental studies, finer-scale trait data, and broader environmental variables to deepen our understanding of the interplay between microbial traits, evolutionary constraints, and environmental filtering.

### Materials and Methods

#### Data sources and homogenization

Data on AM fungal spore traits were obtained from the TraitAM database (70), which compiles species-specific trait information on AM fungi from published taxonomic literature. Spore traits were volume, ornamentation, investment, shape (0 means elongated and 1 means perfect sphere), and color (higher values means higher pigmentation). For AM fungal species within the TraitAM database exhibiting dimorphic spores - glomoid and acaulosporoid types - we focused exclusively on acaulosporoid spores, as their traits were more consistently documented. This selection was applied to *Acaulospora brasiliensis*, *Ambispora appendicula*, *Ambispora fennica*, *Ambispora gerdemannii*, *Ambispora granatensis*, *Ambispora leptoticha*, *Archaeospora ecuadoriana*, and *Archaeospora spainii*. Focusing on this one spore type for dimorphic taxa may weaken any environmental signals present since these taxa could theoretically produce glomoid spores in response to certain environmental pressures but would not lead to false positive signals.

Additionally, we incorporated data on AM fungal presence-absence, abundance, and geographical distribution from the AM fungi biogeography database (30) and the GlobalAMFungi database (31), as compiled by Zahn (71). The AM fungi biogeography database includes species presence-absence data from indexed literature and culture records from the International Collection of (Vesicular) Arbuscular Mycorrhizal Fungi (INVAM), spanning 1922 – 2023. For detailed methodology, please refer to Stürmer et al. (30). Our data on species abundance, target gene, and sequencing platform were sourced exclusively from the GlobalAMFungi database. The GlobalAMFungi database compiles studies from the literature published between 1970 and 2020; details can be found in Větrovský et al. (31). We integrated all three databases (TraitAM, AM fungi biogeography, and GlobalAMFungi) to construct metadata that includes AM fungal spore traits, geographical coordinates (longitude and latitude), biomes, presence-absence, and the estimated abundance of AM fungal species. Biomes were classified as anthropogenic, aquatic, cropland, desert, forest, grassland, shrubland, tundra, and wetland (31).

To examine the relationship between spore traits and climate, we obtained long-term climate variables (1970-2000) from the WorldClim database, version 2.1 (Fick and Hijmans, 2017), at a spatial resolution of approximately 1 km². The climate variables we extracted included mean annual temperature, maximum temperature of the warmest month (maximum temperature), minimum temperature of the coldest month (minimum temperature), temperature seasonality (coefficient of variation of monthly temperature over the year), mean annual precipitation, maximum precipitation of the wettest quarter (maximum precipitation), minimum precipitation of the driest quarter (minimum precipitation), precipitation seasonality (coefficient of variation of monthly precipitation over the year), mean annual wind speed and mean vapor pressure.

We harmonized data across TraitAM, AM fungi biogeography, GlobalAMFungi database, and WorldClim databases. First, trait data from TraitAM were merged with geographical and climatic data from the AM fungi biogeography and GlobalAMFungi databases, matching records by AMF species names. Geographic coordinates from the AM fungi biogeography and GlobalAMFungi databases were then used to extract corresponding climate data from WorldClim. Each ‘site’ was defined as a unique paper reference-latitude-longitude combination. One outlier site with exceptionally high mean annual precipitation was excluded due to unverifiable data. The geographic extent of our dataset, including biomes, is illustrated in Fig. 1.

#### Statistical analyses

All analyses were conducted in R version 4.3.3 (R Core Team, 2023).

### Community-weighted trait-environment relationship

To assess whether AM fungal spore traits vary with climate, we subsetted our metadata to include only sites (N = 1,311) with abundance data and calculated community-weighted mean (CWM) for each trait for each site, weighted by species estimated abundance. We then performed a principal component analysis (PCA) using the ‘princomp’ function in the ‘Hmisc’ package (72) to reduce the dimensionality of all climate variables (mentioned above). Prior to conducting the PCA, all environmental variables were standardized to a mean of zero and a variance of one to ensure comparability. The resulting principal components (PCs) were used as additive predictors in Generalized Additive Models (GAMs) using the ‘mgcv’ package (73). Random effects included biome, the year of sampling, sample type, target gene, and sequencing platform, while spatial autocorrelation was accounted for using a spline-on-the-sphere smooth term based on latitude and longitude (74). Year of sampling was included to control for temporal variability reported in the literature, as environmental conditions and community compositions can fluctuate annually, influencing spore traits. Sample type (field or sediment) was included to account for methodological differences documented in the databases, which could introduce variability unrelated to climatic drivers. Target gene (ITS2, LSU, or SSU) and sequencing platform (454Roche, Illumina, or IonTorrent) were included to control for methodological biases reported in the literature, as differences in gene regions and sequencing technologies can influence detection efficiency and abundance estimates of AM fungal species.

To address deviations from normality, spore volume and shape were Box-Cox transformed, while ornamentation, investment, and color were Yeo-Johnson transformed. The model was initially fitted using restricted maximum likelihood (REML). To account for heteroscedasticity, variance weights were calculated from the initial model and applied in the final GAMs. This framework allowed us to explore the complex interactions between AM fungal spore traits, environmental gradients, and spatiotemporal variability while ensuring robust and reliable parameter estimates. Model diagnostics were performed to evaluate the assumptions of the model, including residual normality, homoscedasticity, and independence (Figure S2).

### Trait-based alpha diversity (Functional diversity)

Trait-based diversity or functional diversity was assessed using three metrics: functional richness, functional evenness (or functional regularity; (75)), and functional divergence. Functional richness reflects the extent to which species occupy functional niche space, representing the cumulative differences among species within an ecological unit (65, 76). This metric relates to the breadth of environmental conditions suitable for species within a community (77). Functional evenness quantifies the consistency of trait differences among species within an ecological unit, where high evenness indicates a more uniform distribution of functional roles across species (65). Functional divergence captures the variance in trait differences among species, reflecting how distinct ecological units are from one another in terms of trait distribution (65).

We calculated functional richness, evenness, and divergence for each site using the ‘fundiversity’ package in R (78). To ensure robust estimates of these functional diversity metrics, only sites with more than five species and available abundance data were included in the analysis (N = 1,258). To examine the relationship between functional diversity and environmental gradients, we fitted GAMs for each of the diversity metrics after log-transformation in the ‘mgcv’ package in R (73). All climate variables (mentioned above) were represented by the first four principal components (PC1 – PC4). Prior to PCA, climate variables were standardized to ensure comparability. The model incorporated random effects to account for variation due to sampling conditions and methodology. Specifically, random effects were included for biome, target gene, sequencing platform, sample type, and the year of sampling (see above). To account for spatial autocorrelation, we included a two-dimensional spline-on-the-sphere smoothing term for latitude and longitude (74). The model was fitted using restricted maximum likelihood (REML) estimation.

### Trait-based beta diversity

To assess how AM fungal spore traits differ between sites and calculate trait-based beta diversity, we subsetted our dataset to include only sites with abundance data and more than five species (N= 378). We then applied Generalized Dissimilarity Modeling (GDM) using the ‘gdm’ package in R (79) to quantify functional beta diversity. GDM is a nonlinear, multivariate modeling approach that identifies the environmental variables best predicting beta diversity and evaluates the extent to which geographical distances between sites explain this variation. By modeling beta diversity as a nonlinear function of environmental gradients, GDM effectively handles complex relationships and accounts for varying rates of beta diversity change along these gradients. To compute the trait-based beta diversity, we first calculated Euclidean distances among traits, referred to as ‘trait distance,’ which served as the response variable in the GDM analysis. This approach allowed us to capture trait-based dissimilarity between pairs of sites as a function of environmental and geographic variation.

### Phylogenetic signal of traits

We analyzed the phylogenetic signal of AM fungal spore traits using a pre-existing phylogenetic tree constructed using LROR-FLR2 fragment of the large ribosomal subunit in the TraitAM database (70), which included 189 AMF species overlapping with our dataset. To visualize trait variation across the phylogeny, we standardized traits to z-scores and constructed a phylogenetic heatmap using the ‘phytools’ package in R (80). In addition, to quantify the phylogenetic signal of traits, we calculated Pagel’s λ using the ‘phylosig’ function in the ‘phytools’ package in R (80). Pagel’s λ measures the extent to which trait variation is explained by phylogeny, ranging from 0 (no phylogenetic signal) to 1 (complete phylogenetic dependence). This analysis provides insights into the degree of evolutionary constraint on each trait and helps identify traits shaped predominantly by phylogenetic history versus those influenced by ecological or environmental factors.

### Trait effect on species range size

Species range size was estimated using the ‘lets.rangesize’ function from the ‘letsR’ package in R (81) as the geographical extent (latitude and longitude) covered by each species. To investigate the relationship between range size and spore traits, we employed a Random Forest model, with range size as the response variable and spore traits (volume, ornamentation, investment, shape, and color) and environmental variables: mean annual temperature (MAT), mean annual precipitation (MAP), soil pH, mean annual wind speed, and precipitation seasonality as predictors. Random Forest modeling was performed using the ‘randomForest’ package (82), which allowed us to account for complex interactions and nonlinear relationships between predictors and range size. Predictor importance was assessed using the mean decrease in accuracy, and partial dependence plots were constructed using “pdp” package in R (83) to visualize individual predictor effects.

## Acknowledgments

The authors would like to thank students/researchers in Chaudhary, Zahn, and Stürmer labs who helped with data collection and cleaning.

