## Supplemental figures for "Climate-linked biogeography of mycorrhizal fungal spore traits"

**Supplementary figures**

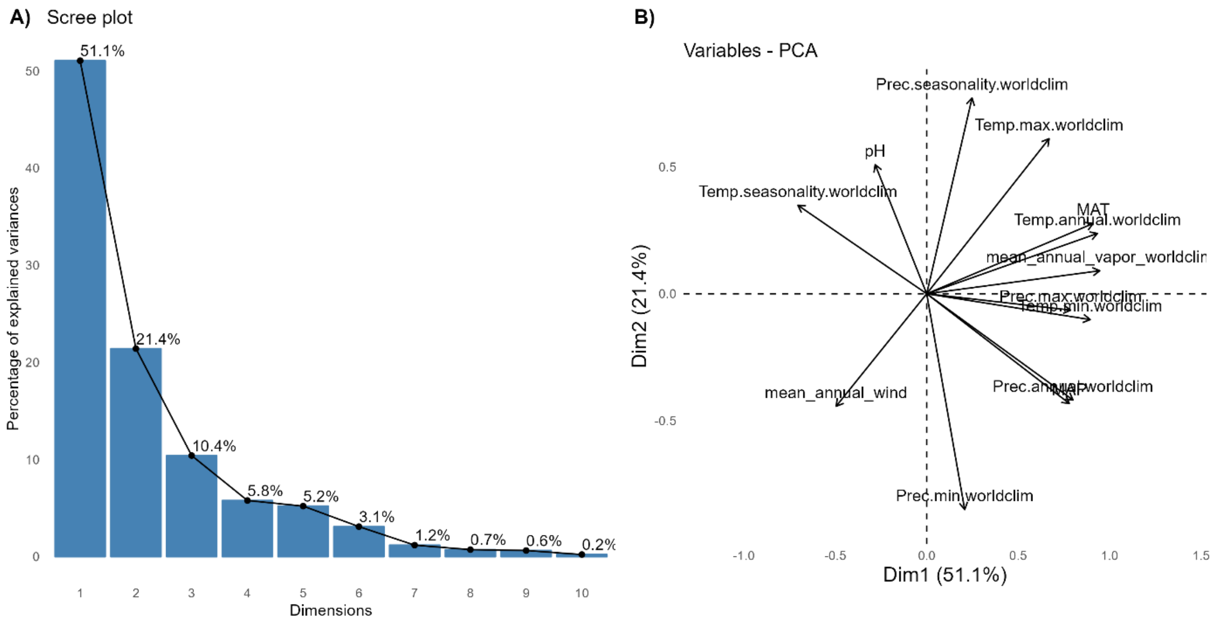


**Figure S1:** Principal Component Analysis (PCA) results for environmental variables used in the analysis of community weighted mean (CWM) trait-environment relationship. (A) Scree plot showing the percentage of explained variance for each principal component (PC). The first four components (PC1, PC2, PC3, and PC4) explain 51.1%, 21.4%, 10.4%, and 5.8% of the variance, respectively, with diminishing returns for the subsequent dimensions. (B) PCA biplot of environmental variables, indicating their contributions to the first two principal components. Variables such as MAT (Mean Annual Temperature), Temp.min (Minimum Temperature), and Prec.max (Maximum Precipitation) load heavily on PC1, while variables like Prec.seasonality (coefficent of variation of monthly precipitation over the year) contribute significantly to PC2.

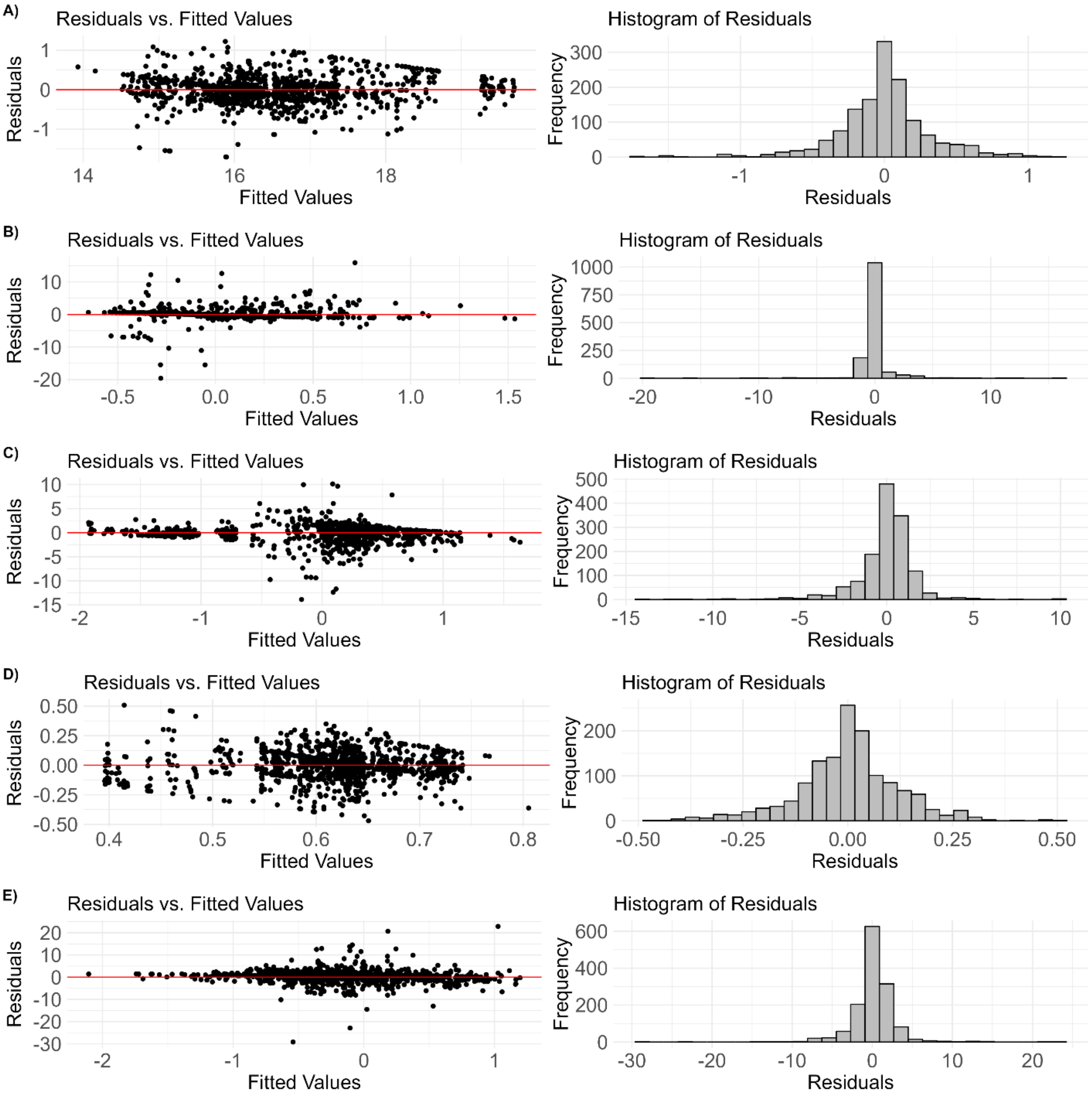


**Figure S2:** Diagnostic plots for generalized additive models (GAMs) evaluating community-weighted mean (CWM) trait-environment relationships across arbuscular mycorrhizal (AM) fungal spore traits: A) Volume, (B) Ornamentation, (C) Investment, (D) Shape, and (E) Color. For each trait, the left panel shows the residuals versus fitted values plot, assessing homoscedasticity and linearity, with a red line at zero indicating no systematic deviation. The right panel displays histograms of residuals, assessing the normality of residuals for each model.

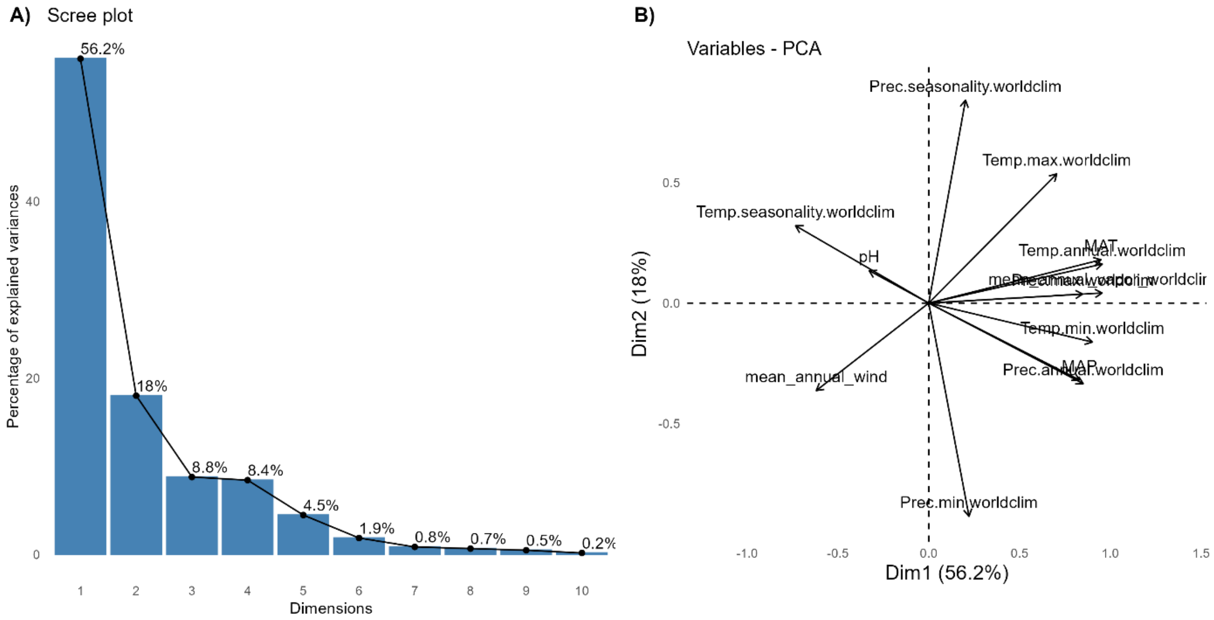


**Figure S3:** Principal Component Analysis (PCA) results for environmental variables used in the analysis of trait-based alpha diversity or functional alpha diversity. (A) Scree plot showing the percentage of explained variance for each principal component (PC). The first four components (PC1, PC2, PC3, and PC4) explain 56.2%, 18%, 8.7%, and 8.4% of the variance, respectively, with diminishing returns for the subsequent dimensions. (B) PCA biplot of environmental variables, indicating their contributions to the first two principal components. Variables such as MAT (Mean Annual Temperature), Temp.min (Minimum Temperature), and Prec.max (Maximum Precipitation) load heavily on PC1, while variables like Prec.seasonality (coefficent of variation of monthly precipitation over the year) contribute significantly to PC2.

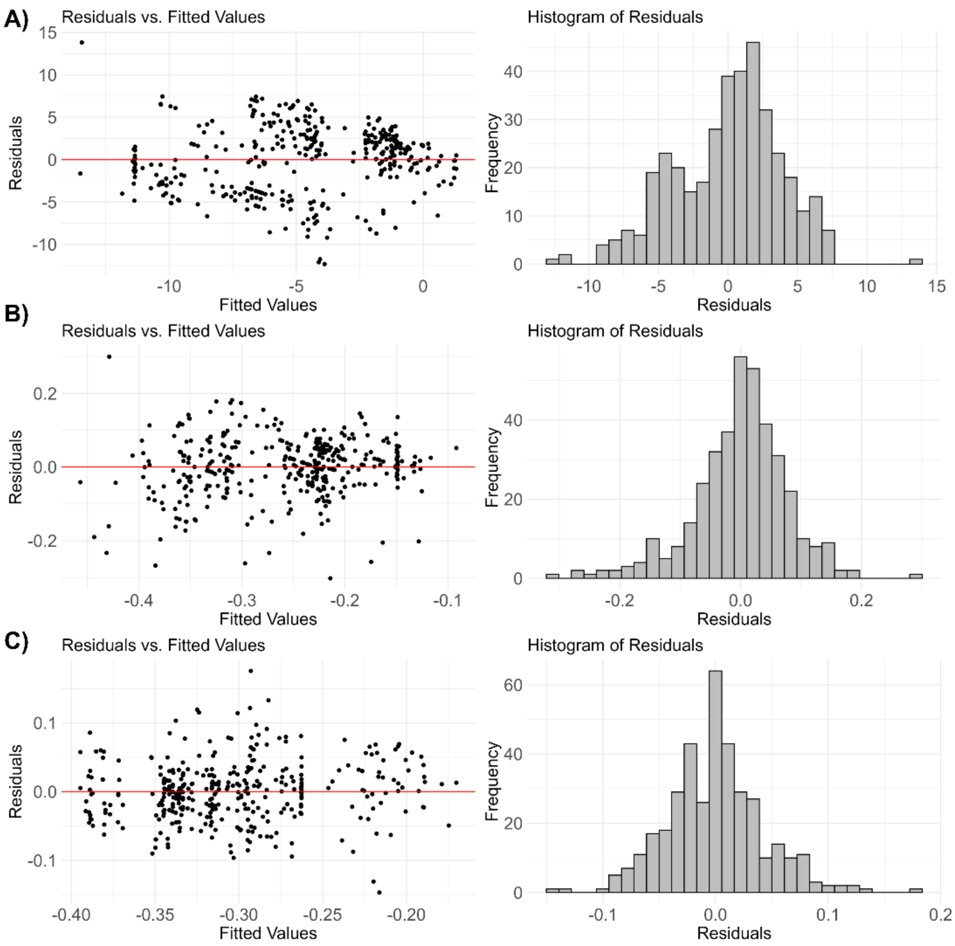


**Figure S4:** Diagnostic plots for generalized additive models (GAMs) evaluating functional alpha diversity across environmental gradients. Each row represents model diagnostics for different functional diversity indices: (A) Functional Richness, (B) Functional Evenness, and (C) Functional Divergence. For each index, the left panel shows the residuals versus fitted values plot, assessing homoscedasticity and linearity, with a red line at zero indicating no systematic deviation. The right panel displays histograms of residuals, assessing the normality of residuals for each model.

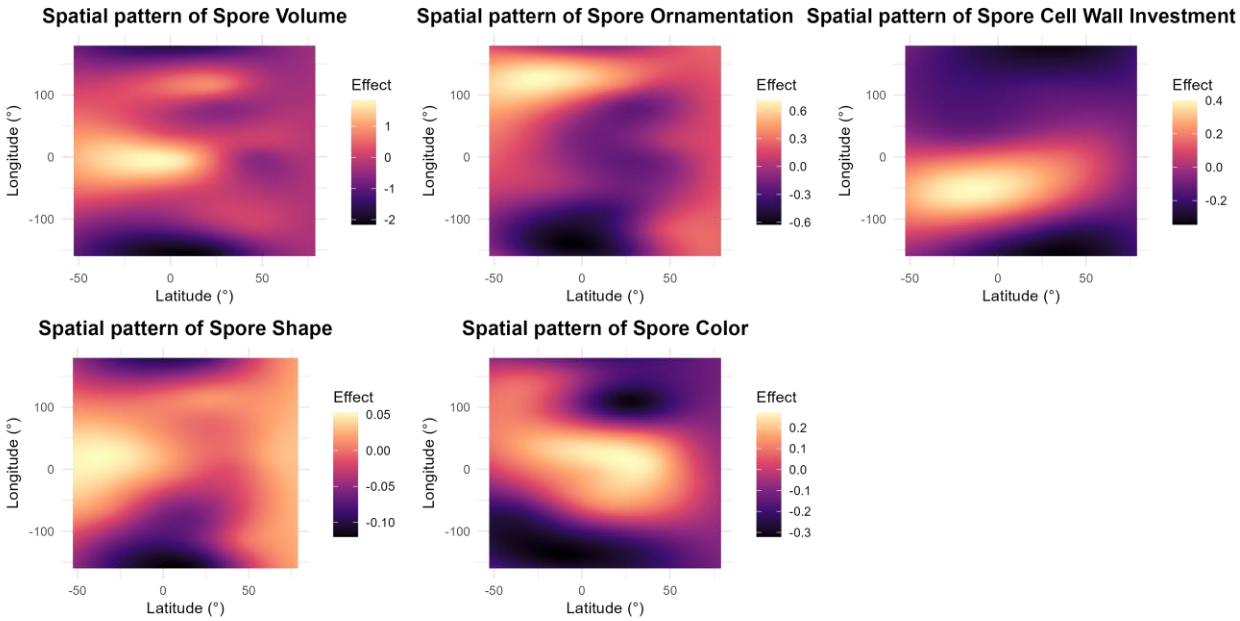


**Figure S5:** Spatial patterns of AM fungal spore traits across latitude and longitude. The plots illustrate the variation in spore volume, ornamentation, cell wall investment, shape, and color along geographic gradients. The color scale indicates the magnitude of the spatial patterns' influence on trait values, with darker hues corresponding to lower values and lighter hues representing higher values.

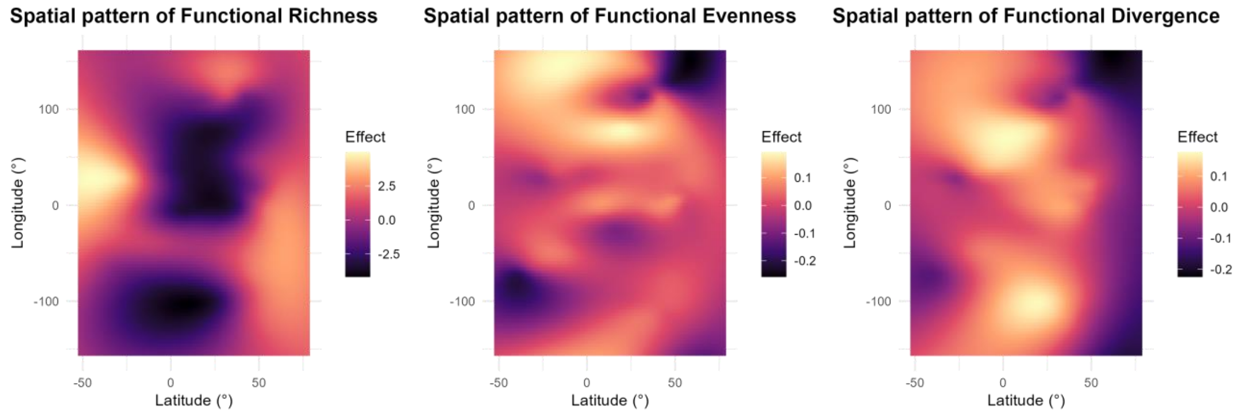


**Figure S6:** Spatial patterns of AM fungal spore traits across latitude and longitude. The plots illustrate the variation in functional richness, evenness, and divergence along geographic gradients. The color scale indicates the magnitude of the spatial patterns' influence on functional diversity values, with darker hues corresponding to lower values and lighter hues representing higher values.
